# IcyTree: Rapid browser-based visualization for phylogenetic trees and networks

**DOI:** 10.1101/110213

**Authors:** Timothy G. Vaughan

## Abstract

**Summary:** IcyTree is an easy-to-use application which can be used to visualize a wide variety of phylogenetic trees and networks. While numerous phylogenetic tree viewers exist already, IcyTree distinguishes itself by being a purely online tool, having a responsive user interface, supporting phylogenetic networks (ancestral recombination graphs in particular), and efficiently drawing trees that include information such as ancestral locations or trait values. IcyTree also provides intuitive panning and zooming utilities that make exploring large phylogenetic trees of many thousands of taxa feasible.

**Availability and Implementation:** IcyTree is a web application and can be accessed directly at http://tgvaughan.github.com/icytree. Currently-supported web browsers include Mozilla Firefox and Google Chrome. IcyTree is written entirely in client-side JavaScript (no plugin required) and, once loaded, does not require network access to run. IcyTree is free software, and the source code is made available at http://github.com/tgvaughan/icytree under version 3 of the GNU General Public License.

## 1 Introduction

The visualization of phylogenetic trees is an extremely important aspect of computational phylogenetics. Indeed, the well-known text Inferring Phylogenies [4] devotes an entire chapter to this topic. It is therefore no coincidence that the evolution of phylogenetic tree visualization software has often gone hand-in-hand with the development of sophisticated algorithms and software for phylogenetic inference. Many of the earliest available phylogenetic inference packages such as PHYLIP [3], MacClade [9], and MrBayes [10] include built-in support for visualizing their output. Other packages such as PhyML [5] and BEAST [1] produce no graphical output but instead rely on specialized phylogenetic visualization tools such as FigTree (http://tree.bio.ed.ac.uk/software/figtree) and Dendroscope [7] to interpret inference results.

There now exists a range of stand-alone phylogenetic visualization tools, too many to name here. (The PHYLIP home page provides an online list of such programs which currently contains 60 entries.) Despite existing solutions to the visualization problem being so numerous, the package IcyTree distinguishes itself in a number of important respects.

Firstly, IcyTree is extremely easy to access. It is written entirely in JavaScript and runs in modern web browsers (compatibility with recent versions Google Chrome and Mozilla Firefox is assured) without needing additional plugins, meaning that it is loaded simply by navigating to its home page. This also means that it is never necessary to update IcyTree. Importantly, and unlike many of the existing JavaScript-based phylogenetics tools, IcyTree does not interact with servers in any way while running. This ensures that the program is as responsive as possible and means that users need not be concerned about the privacy implications of uploading data to an online service.

Secondly, the use of JavaScript combined with the SVG graphical format allows IcyTree to display trees containing thousands or even tens of thousands of taxa while remaining responsive. This also means that large files containing thousands of trees can be quickly loaded.

Thirdly, IcyTree is very light-weight, focusing on drawing rooted phylogenetic time trees such as those inferred by Bayesian phylogenetic inference packages. This keeps the number of available options manageable. Along with the emphasis placed on making the program options accessible using a mouse or via an extensive repertoire of keyboard shortcuts, I feel that this makes IcyTree intuitive and easy to use, catering to new and seasoned users alike.

Finally, IcyTree is capable of viewing trees stored in a number of different formats, including Newick, NEXUS [8] (as used by BEAST and MrBayes), PhyloXML [6] and NeXML [13]. Phylogenetic networks specified using the Extended Newick format [2] are also supported. (The style used by IcyTree to render such networks is particularly suited to ancestral recombination graphs such as those inferred by Bacter [12].)

**Fig. 1.**
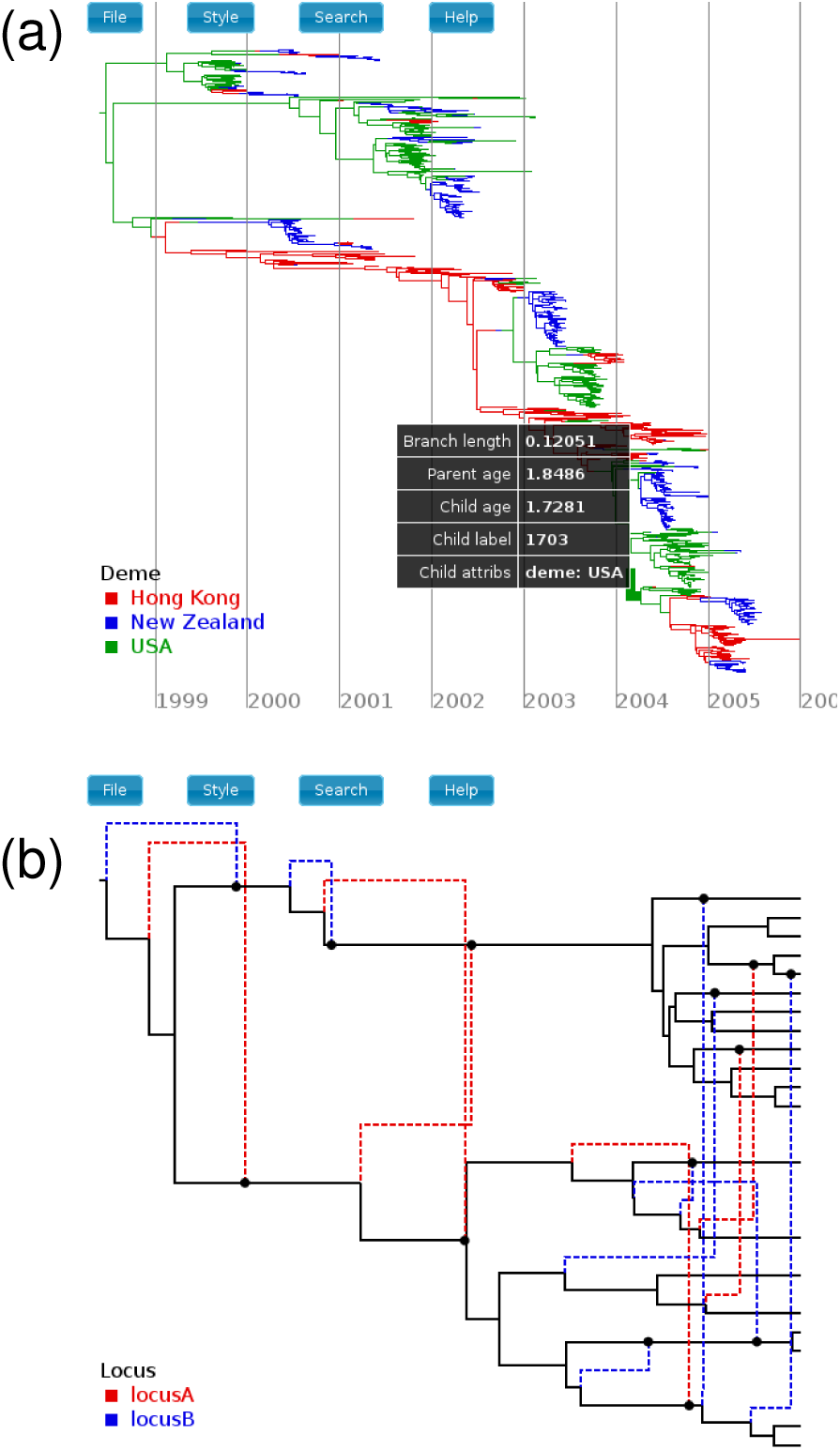
(a) Rendering of an influenza tree annotated by edge locations (colours). The table lists information specific to a particular edge selected using the mouse. (b) Rendering of a phylogenetic network loaded from its Extended Newick description. Dashed edges represent reticulations, while the colour shows the values of an annotation representing the locus of a recombination event.

## 2 Basic use and capabilities

IcyTree is opened by navigating to its URL http://tgvaughan.github.io/icytree using either Mozilla Firefox or Google Chrome. (Chrome runs IcyTree noticeably faster than Firefox, but both browsers are supported.) Tree files can then be loaded by simply dragging and dropping them onto the IcyTree window, or by selecting File→Open. (Newick strings can also be pasted directly into IcyTree by using File→Enter tree directly.)

Once a tree file is displayed, it can be easily explored using the pan-and-zoom interface: the mouse scroll wheel is used to zoom in or out of features under the mouse cursor, while the displayed area can be panned by clicking and dragging the mouse. (This is a similar interface to that provided by online map services.) In the instance that tree files contain multiple trees, the displayed tree can be changed using either the navigation box that appears in the bottom-left corner of the screen or via keyboard shortcuts.

Moving the mouse cursor over a tree edge displays a table containing information about that edge and the nodes that it connects. This includes the label of the child node, the age of the child and parent nodes, the length of the edge and the values of any additional annotations.

The Style menu contains a number of options which affect how the tree is displayed, such as changing what text is displayed on the tree, which annotations if any determine edge colour or opacity, whether reticulations are displayed, axis options, etc.

Figures produced by IcyTree can be exported directly as scalable SVG images via the File→Export→SVG menu item. Bitmap images can also be produced using the adjacent PNG and JPEG menu items.

All of IcyTree’s features are described in an extensive usage manual which is available via the Help→Online Manual menu item.

## 3 Supported input formats

IcyTree can read tree files produced by popular phylogenetics packages such as MrBayes, BEAST and PhyML. These programs produce files in Newick and NEXUS formats, with the NEXUS format supporting the BEAST extensions which allow metadata (e.g. clade posterior support, ancestral traits, etc.) to be associated with each node/edge in a tree. The support for Extended Newick is built into IcyTree at the lowest level, and allows IcyTree to not only read plain network descriptions but also NEXUS tree blocks containing annotated phylogenetic networks. These annotations are used by programs such as MASTER [11] and Bacter [12] to represent ancestral trait information.

In addition, IcyTree can be used to view trees represented using the modern PhyloXML and NeXML formats, although support for these formats is currently limited to trees only.

## References

[1] R. Bouckaert et al. BEAST 2: a software platform for Bayesian evolutionary analysis. PLoS Comput. Bio., 10(4):e1003537, Apr 2014.

[2] G. Cardona et al. Extended Newick: it is time for a standard representation of phylogenetic networks. BMC Bioinf., 9:532,2008.

[3] J. Felsenstein. Phylip - phylogeny inference package (version 3.2). Cladistic., 5:164–166,1989.

[4] J. Felsenstein. Inferring Phylogenies. Sinauer Associates, Massachusetts, 2003.

[5] S. Guindon and O. Gascuel. A simple, fast, and accurate algorithm to estimate large phylogenies by maximum likelihood. Syst. Biol., 52(5):696–704, Oct 2003.

[6] M.V. Han and C. M. Zmasek. phyloXML: XML for evolutionary biology and comparative genomics. BMC Bioin., 10:356,2009.

[7] D. H. Huson and C. Scornavacca. Dendroscope 3: an interactive tool for rooted phylogenetic trees and networks. Syst. Biol., 61(6):1061–1067, Dec 2012.

[8] D. R. Maddison et al. NEXUS: an extensible file format for systematic information. Syst. Biol., 46:590–621, Dec.1997.

[9] W. P. Maddison and D. R. Maddison. Interactive analysis of phylogeny and character evolution using the computer program macclade. Folia Primatol (Basel), 53(1–4):190–202,1989.

[10] F. Ronquist et al. MrBayes 3.2: efficient bayesian phylogenetic inference and model choice across a large model space. Syst. Bio., 61(3):539–542, May 2012.

[11] T. G. Vaughan and A. J. Drummond. A stochastic simulator of birth-death master equations with application to phylodynamics. Mol. Biol. Evol., 30(6):1480–1493, Jun 2013.

[12] T. G. Vaughan et al. Inferring ancestral recombination graphs from bacterial genomic data. Genetic., 205:857–870, Feb.2017.

[13] R. A. Vos et al. NeXML: rich, extensible, and verifiable representation of comparative data and metadata. Syst. Biol., 61(4):675–689, Jul 2012.

